# The AWSEM-Amylometer: predicting amyloid propensity and fibril topology using an optimized folding landscape model

**DOI:** 10.1101/138842

**Authors:** Mingchen Chen, Nicholas P Schafer, Weihua Zheng, Peter G Wolynes

## Abstract

Amyloids are fibrillar protein aggregates with simple repeated structural motifs in their cores, usually *β*-strands but sometimes *α*-helices. Identifying the amyloid-prone regions within protein sequences is important both for understanding the mechanisms of amyloid-associated diseases and for understanding functional amyloids. Based on the crystal structures of seven cross-*β* amyloidogenic peptides with different topologies and one recently solved cross-*α* fiber structure, we have developed a computational approach for identifying amyloidogenic segments in protein sequences using the Associative memory, Water mediated, Structure and Energy Model. The AWSEM-Amylometer performs favorably in comparison with other predictors in predicting aggregation-prone sequences in multiple datasets. The method also predicts the specific topologies (the relative arrangement of *β*-strands in the core) of the amyloid fibrils well. An important advantage of the AWSEM-Amylometer over other existing methods is its direct connection with an efficient, optimized protein folding simulation model, AWSEM. This connection allows one to combine efficient and accurate search of protein sequences for amyloidogenic segments with the detailed study of the thermodynamic and kinetic roles that these segments play in folding and aggregation in the context of the entire protein sequence. We present new simulation results that highlight the free energy landscapes of peptides that can take on multiple fibril topologies. We also demonstrate how the Amylometer methodology can be straightforwardly extended to the study of functional amyloids that have the recently discovered cross-*α* fibril architecture.

## Introduction

Amyloid formation by proteins and peptides has been the focus of a tremendous amount of research.^1, 2^ A large and growing body of evidence suggests that amyloid formation plays a role both in functional^3^ and in pathological biological processes.^4^ The amyloid fibril based on *β*-strands is a common, though not universal, aggregate architecture. The propensity of a full-length protein to form amyloid has been linked to the presence of short sequences, typically five to seven residues in length, within longer protein sequences. In isolation, these short “amyloidogenic” segments by themselves oftentimes readily form fibrils, and, therefore, many *in vitro* studies have focused on these short peptides. ^5^ Some natural peptides that form amyloids *in vivo* are indeed short and largely disordered as monomers, such as A*β* and *α*-synuclein,^6^ though at 40 to 140 residues in length these protein fragments are still long compared to those parts of the sequences that seem to be primarily responsible for initiating aggregation.

In the case of amyloid-forming proteins that also fold to a native structure, an even larger proportion of the sequence lies outside of the amyloidogenic segment that eventually makes its way into amyloid fibril cores when the balance between folding and aggregation is upset by, e.g., a destabilizing mutation, high protein concentration, high temperatures, or a change in solvent conditions. The entire sequence, including the parts that apparently never become incorporated into the fibril core, can play a role throughout the aggregation process. Folding to the native state, in general, is the result of cooperation between a diffuse but structurally consistent set of stabilizing interactions throughout the folded structure. ^7^ These ‘minimally frustrated’ interactions predominate in the core of natively folded protein structures, which is also where the most amyloidogenic segments within a protein sequence typically are buried. ^8^ When a protein unfolds and starts to form oligomers, parts of the sequence outside of the primary amyloidogenic segment influence the size, shape, and stability of the oligomers. ^9^ Finally, unless extensive proteolytic processing precedes fibril formation, the entire sequence must also be accommodated in the mature aggregates and disordered parts of the structure thus may make important entropic contributions to the stability.

In protein aggregation, multiple copies of amyloidogenic segments in close proximity can recognize each other and become stabilized in a misfolded/aggregated state. ^10^ The self-recognition of protein domains in repeat proteins with high identity in sequence has been extensively studied by Jane Clarke and her coworkers.^11, 12^ While domain swapping is a major contributor to misfolding, simulation studies revealed that *I*27 domains from titin initially aggregate by means of an amyloidogenic segment which has a strong tendency to self-recognize. ^10, 11^ In contrast, *SH*3 dimers, which do not possess any amyloidogenic segments, don't aggregate significantly. ^10, 11^

In this context, the identification of amyloidogenic segments within protein sequences using local information alone can be only a first step in elucidating amyloid formation. Most existing models and algorithms for identifying these segments,^5, 13–19^ however, are poorly suited for following on to address the mechanistic questions that arise naturally once an amyloidogenic segment has been identified within a protein sequence. How does a given segment contribute to folding, misfolding, oligomerization, and aggregation? And how does the rest of the sequence affect these same processes? At the same time, even with recent advances in computer algorithms and hardware, addressing such questions using fully atomistic models remains difficult. To overcome this difficulty, here we introduce a method for detecting amyloidogenic segments that is based on the Associative Memory, Water mediated, Structure and Energy Model (AWSEM), an optimized, coarse-grained, protein folding simulation model. ^20^ AWSEM has been fruitfully applied in recent years to many different problems of protein structure prediction, ^20–22^ protein association, ^23^ allosteric mechanism ^24^ and protein aggregation. ^25–29^ The AWSEM-Amylometer is based on the same energy model that is used in AWSEM molecular dynamics simulations but is able to detect amyloidogenic segments using a simple and efficient threading scheme over multiple fiber template structures. This scheme not only allows for the detection of amyloidogenic segments but also the prediction of the relative orientation of the amyloid *β*-strands in the fibril core. Moreover, the efficiency of the AWSEM-Amylometer and its connection to AWSEM allows surveys of large numbers of protein sequences, including naturally occurring and designed mutants to be accompanied by selective followup studies using statistical and structural analyses of dynamic simulations. In the following sections we introduce the AWSEM-Amylometer scanning methodology and discuss its prediction accuracy when tested on databases of peptides and proteins. We also present some new simulations on amyloidogenic peptide aggregation, and extend the methodology to the study of amyloids with the recently discovered cross-*α* functional fibril architecture.

## Method

### 1: The AWSEM force field

AWSEM (the Associative Memory, Water Mediated, Structure and Energy Model) is a predictive, coarse-grained, protein folding force field that represents amino acids using three explicit interaction sites per residue. AWSEM's parameters were optimized using a database of solved protein structures and the principles of energy landscape theory. ^7^ Interested readers are encouraged to consult Davtyan et al. ^20^ for detailed information about the AWSEM force field. The AWSEM Hamiltonian is summarized in Eq. 1. AWSEM includes a fragment-based associative memory term, *V_FM_*, that locally biases the formation of secondary and super-secondary structures. This bias can be based on using experimentally solved structures from the Protein Data Bank (PDB) with or without knowledge of global sequence homology as input. Alternatively this bias can employ structures sampled in atomistic simulations.^21, 22^ The backbone term, *V_backbone_*, ensures that the peptide backbone stays connected and does not overlap itself. The many body burial term, *V_burial_*, takes into account the instantaneous local density around each residue and attempts to sort each residue into its preferred burial environment-exposed, partially buried, or completely buried. The contact term, *V_contact_*, includes a direct contact interaction and a water- or protein-mediated interaction. The hydrogen bonding term, *V_HB_*, favors formation of *α*-helices or *β*-sheets.

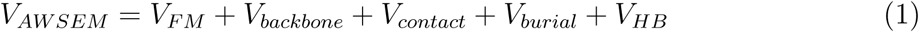

The AWSEM-Amylometer energy function (Eq. 2) used to detect amyloidogenic segments using threading over template fiber structures does not include the associative memory term, *V_FM_*, because evaluation of *V_FM_* requires either homology searches or atomistic simulations to be performed before carrying out further calculation. These steps would be incompatible with a rapid threading scheme like that which the AWSEM-Amylometer uses to predict amyloid propensity. At the same time, amyloid structures are apparently under-represented among the existing solved structures in the PDB considering how common the amyloid architecture seems to be. Only 104 fiber structures have been solved to date. Thus, structural constraints from known structures would artificially disfavor amyloid-compatible conformations. In the amylometer, secondary structure preferences are thus accounted for solely by the hydrogen bonding term, *V_HB_*. The backbone term, *V_backbone_*, is sequence independent and therefore is also left out of the AWSEM-Amylometer calculations.

The AWSEM-Amylometer works by first threading protein sequences, typically six residues at a time, over experimentally determined fiber structures and then evaluating the potential energy of each of those candidate structures. In its simplest instantiation, AWSEM-Amylometer-Min, a protein sequence segment will be considered to be highly aggregation/amyloid prone if the energy of that segment in any of the cross-*β* fibril structures is below an empirically determined threshold value (−100 kcal/mol). ^10^ Consideration of the propensity to form cross-*α* fibers is done separately and will be discussed in Section 6.

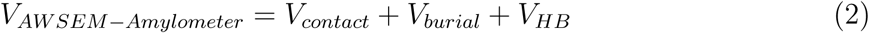

### 2: Predicting fibril topology using multiple fiber templates

The idea of a direct structure-based approach to prediction of fibril formation was introduced by Eisenberg and coworkers. They used the crystal structure of the fibril-forming hexapeptide NNQQNY (PDB ID: 1YJO) from sup35 prion protein from yeast as a template. ^14^ They were able to show that threading protein sequences onto this template or a template ensemble derived from crystal structures could yield reasonably accurate predictions of amyloidogenic regions. Since the publication of the 3D-Profile method, many more fiber structures, including an *α*-helical fiber, have been solved. The AWSEM-Amylometer takes advantage of these multiple fiber structures to predict not only amyloid propensity but also to predict specific fibril topology.

Cross-*β* fibril structures can be classified into 8 classes based on the relative orientation of the *β*-strands within the *β*-sheets (parallel or antiparallel) and the relative orientations of the *β*-sheets that are further packed together (Fig. 1). A total of 24 hexapeptide crystal structures, which cover seven of the eight classes (class 3 is missing), are currently available. We chose 7 structures, one from each available class, over which to thread hexapeptide sequences (Figure 1). The energy of a hexapeptide is evaluated on each of the seven templates separately, and the class corresponding to the template with the lowest energy is then the predicted fibril topology. To predict whether a hexapeptide will form parallel or anti-parallel sheets, the lowest score among the parallel cross-*β* spines (classes 1, 2 and 4) is compared with the lowest score from the anti-parallel cross-*β* spines (classes 5, 6, 7 and 8), and the hexapeptide is predicted to have the orientation corresponding to the template with the lowest energy. For testing the possibility of favoring an *α*-helical amyloid, we used the recently determined fiber structure of PSMα3 (22 residues, PDB ID: 5I55) as the template (Figure 1). The threshold for predicting that a 22-residue peptide will assume this fibril structure was determined based on the statistics of 5000 random sequences such that only 5% of the sequences gave energy values below this threshold. The corresponding threshold value is −205kcal/mol.

**Figure 1:**
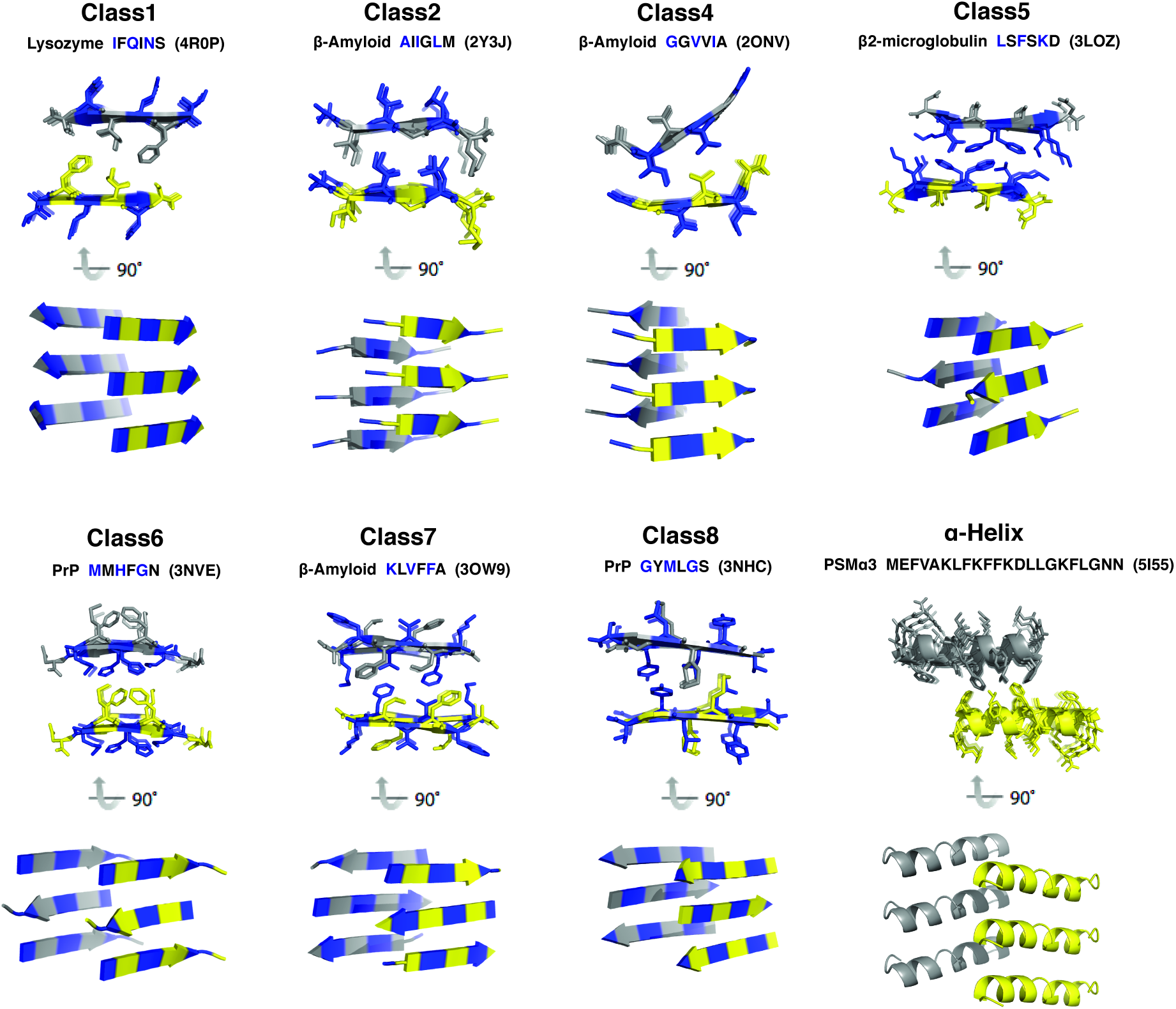
Templates for the 7 cross-*β* and one cross-*α* classes used by the AWSEM-Amylometer, with views both parallel to the fibril axes (the interdigitation of side chains is shown) and perpendicular to the fibril axes. The name of the protein that the peptide is derived from, the sequence of the peptide, and the PDB ID of the template structure are given above each class. Class 1, class 2 and class 4 structures have parallel, in-register *β*-sheets, while class 5 to class 8 have anti-parallel *β*-sheets. The seven types of steric zippers are organized into symmetry classes depending on the relative orientations of the two *β*-sheets the *β*-strands within the *β*-sheets. Different sheets are shown in different colors (yellow and gray). The first, third and fifth residues of the *β*-strands are colored blue to clarify the different orientations of the sheets. The cross-*α* template contains *α*-helices only. Abbreviations: PrP, prion protein; PSM*α*3, phenol-soluble modulin *α*3.

### 3: Metrics used to evaluate prediction capacity of the AWSEM-Amylometer

To evaluate and compare the performances of different predictors, we used the following five classical quantitative evaluation measures: accuracy (Eq. 3), sensitivity (Eq. 4), specificity (Eq. 5), F1 score (Eq. 6) and Matthews Correlation Coefficient (MCC, Eq. 7).

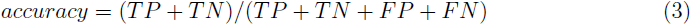

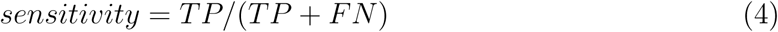

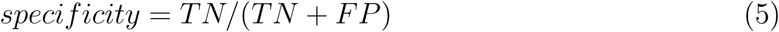

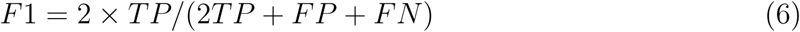

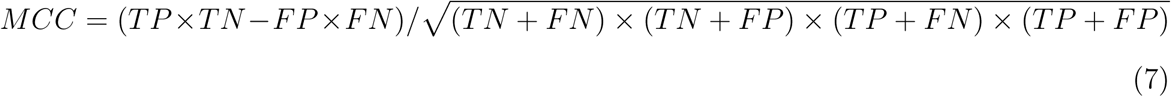

In Eqs. 3–7, *TP* is the number of true positives predicted by the algorithm, *TN* is the number of true negatives, *FP* is the number of false positives, and *FN* is the number of false negatives.

Using these evaluation measures, we compared the performance of AWSEM-Amylometer with the performance characteristics of several other amyloid predictors including the 3D profile method, ^14^ AGGRESCAN, ^15^ FoldAmyloid, ^18^ PAFIG, ^17^ PASTA, ^19^ SALSA, ^16^ TANGO ^13^ and Waltz ^5^ using the amylome dataset. ^30^ We also compared the performance of AWSEM-Amylometer with that of TANGO and Waltz on the Waltz dataset.

### 4: Parameter fitting using linear regression for propensity to form cross-*β* structures

For cross-*β* propensity predictions, the seven different *β*-spine topologies generate seven different predicted scores. In its simplest version, AWSEM-Amylometer-Min, the algorithm merely checks whether any of these scores is below a threshold. Apart from the threshold, the parameters in AWSEM-Amylometer-Min are all obtained from the AWSEM force field itself. It is possible to improve somewhat the prediction power of the algorithm by training a composite model wherein all 7 individual predictors are weighted differently by tuning 8 coefficients in Eqs. 8. The following enhanced linear regression model yields a composite score, *f* (*sequence*), that is the score used to predict whether a hexapeptide is amyloid/aggregation prone. *E_ClassN_*, *N* = 1 − 2, 4 − 7 are the amyloidogenic energies on 7 cross-*β* templates. The optimized values of the regression coefficients (available in SI) is achieved by maximizing the likelihood of a logistic model.

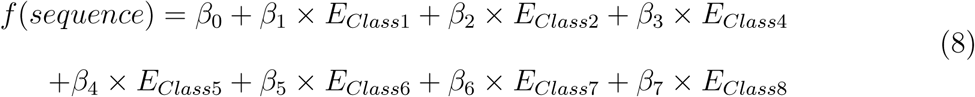

To optimize these fine-tuning parameters, as well as the threshold for cross-*β* amyloid formation, we split the Waltz dataset (1088 hexa-peptides) into a training set (816 hexa-peptides) and a test set (272 hexa-peptides). We carried out linear regression on the training dataset and used the trained parameters to examine the test set, and the cutoff value was selected as the point with the highest Matthews Correlation Coefficient (MCC) on the test set. The threshold score, *f*, of the hexapeptide to be predicted amyloidogenic, after linear regression of the seven input energy values, is 0.5.

The simpler model, AWSEM-Amylometer-Min, is more directly physical and uses only the minimal value of the seven individual predictors. Its predictions are also compared in the result section. In this approach, if the minimal value out of the seven predictors for a given hexapeptide is below the determined threshold (−100kcal/mol), ^10^ this hexapeptide will be considered to be amyloid-prone.

### 5: Simulation details using the physics-based AWSEM force field

Detailed molecular dynamics simulations of peptide aggregation for some examples were performed using the Large-Scale Atomic/Molecular Massively Parallel Simulator (LAMMPS) software package, in which the AWSEM force field is available in open source format. ^20^ All the umbrella sampling simulations for multiple peptide chains were performed at 300K for 20 million steps. 20 million steps corresponds to roughly 0.1 ms in laboratory time in the AWSEM force field. At the simulation concentration, this time is long enough to ensure the convergence of sampling on this system when using umbrella sampling. The initial configurations used for the umbrella sampling were ten monomers randomly distributed over a cubic box of size 100 Angstroms.

### 6: Order parameter for umbrella sampling and free energy calculations

To compute the relative free energy of forming parallel versus anti-parallel topologies for a set of 10 hexapeptides, we used umbrella sampling along an order parameter, *Q_diff_* (Eq. 10), to sample structures both near the limits and intermediate between the two topologies.

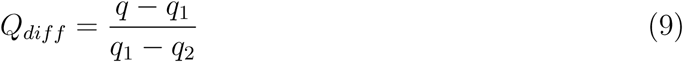

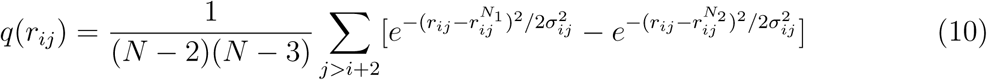

In Eq. 10, 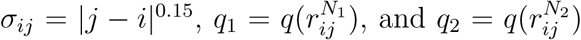 where the superscripts *N*_1_ and *N*_2_ indicate distances evaluated in the anti-parallel and parallel fibril structures.

The harmonic potential used for constant temperature umbrella sampling simulations along *Q_diff_* is shown in Eq 11.

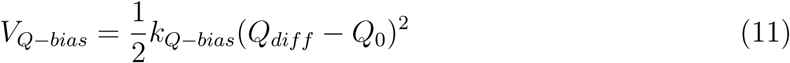

In Eq. 11, *k_Q−bias_* = 200*kcal/mol*. The biasing center values *Q*_0_ were chosen to be equally spaced from 0 to 0.98 with a step size 0.02. The unbiased free energy landscapes were then reconstructed from the umbrella sampling data using the weighted histogram analysis method (WHAM).^31^

## Results and discussion

### 1: Performance of the AWSEM-Amylometer on the Waltz peptide dataset for predicting cross-*β* amyloid propensity

To test the ability of the AWSEM-Amylometer to predict the propensity of hexapeptides to form cross-*β* amyloid of any topology, we examined the performance of AWSEM-Amylometer-Min and the complete AWSEM-Amylometer based on a composite score using a subset of the Waltz dataset (details in Methods). This dataset contains experimental information about amyloid formation for 1088 hexa-peptides. The composite linear regression model of the threading energies was obtained with optimized coefficients from a training subset of the Waltz dataset (716 hexa-peptides) and a threshold score (0.5) was determined based on a validation subset from the Waltz dataset (272 hexa-peptides). When we applied this fully optimized model to the whole dataset, in terms of accuracy, the full AWSEM-Amylometer outperformed the other methods with a correct classification rate of 0.84 (Table 1). To quantify the advantage of the composite model over using only a single topology, we compared the prediction performances of several variants of the AWSEM-Amylometer: one variant using the combined score from the linear regression model, one taking only the minimum score from the seven topologies, and one using the scores of the class 1 topology and class 8 topology alone. The complete AWSEM-Amylometer using the composite linear model with optimized parameters has a prediction performance somewhat higher than AWSEM-Amylometer-Min. Predictions from Waltz and TANGO have accuracies of 0.77 and 0.80 respectively, which are lower than the complete AWSEM-Amylometer models and comparable to AWSEM-Amylometer-Min (Table 1).

**Table 1:** Evaluation of the performance on the training dataset.

| Predictor | TP | TN | FP | FN | Accuracy (%) | Sensitivity (%) | Specificity (%) | MCC | F1 |
| --- | --- | --- | --- | --- | --- | --- | --- | --- | --- |
| **AWSEM-Amylometer** | 151 | 760 | 92 | 85 | 83.73 | 62.14 | 89.94 | 0.53 | 0.63 |
| **AWSEM-Amylometer (Min)** | 164 | 687 | 158 | 79 | 78.22 | 67.49 | 81.30 | 0.45 | 0.58 |
| **AWSEM-Amylometer (Class1)** | 139 | 752 | 93 | 104 | 82.81 | 57.20 | 88.99 | 0.47 | 0.59 |
| **AWSEM-Amylometer (Class8)** | 148 | 755 | 90 | 95 | 82.99 | 60.91 | 89.35 | 0.51 | 0.61 |
| **Waltz** | 166 | 668 | 177 | 77 | 76.65 | 68.31 | 79.05 | 0.42 | 0.57 |
| **TANGO** | 59 | 816 | 29 | 184 | 80.42 | 24.28 | 96.57 | 0.32 | 0.36 |

When we examine other evaluation measures, we find the TANGO method has high specificity but low sensitivity. The AWSEM-Amylometer with a single topology (Class 1 or 8) also sometimes fails to recognize an amyloidogenic segment, perhaps because those peptides prefer a different topology. Not surprisingly, the AWSEM-Amylometer using the minimum score across all topologies achieves a somewhat higher sensitivity but a lower specificity. In comparison, the AWSEM-Amylometer using the regression score has a more balanced specificity and sensitivity.

### 2: Performance of the AWSEM-Amylometer on predicting the amyloidogenic regions in complete proteins found from an amylome dataset

An important application of a predictor such as the AWSEM-Amylometer is to identify the primary fibril-forming segments within full-length proteins so as to provide predictions that can be useful for guiding experimental studies on natural proteins. We compared the AWSEM-Amylometer with 9 other tools for detecting amyloid-prone regions in a set of 33 proteins belonging to the amylome. ^30^ This test set was constructed by Tsolis et al who searched to find data from many published experiments and different experimental methods that support the amyloidogenicity of specific regions in the 33 proteins of the set. ^30^ In terms of predicting the amyloidogenic regions in these 33 long sequences, the AWSEM-Amylometer performs well as judged by the MCC and F1 scores (Table 2). Only PAFIG (0.18) and AMYLPRED2 (0.20) yield slightly higher MCC scores than does the AWSEM-Amylometer (0.17).

**Table 2:**
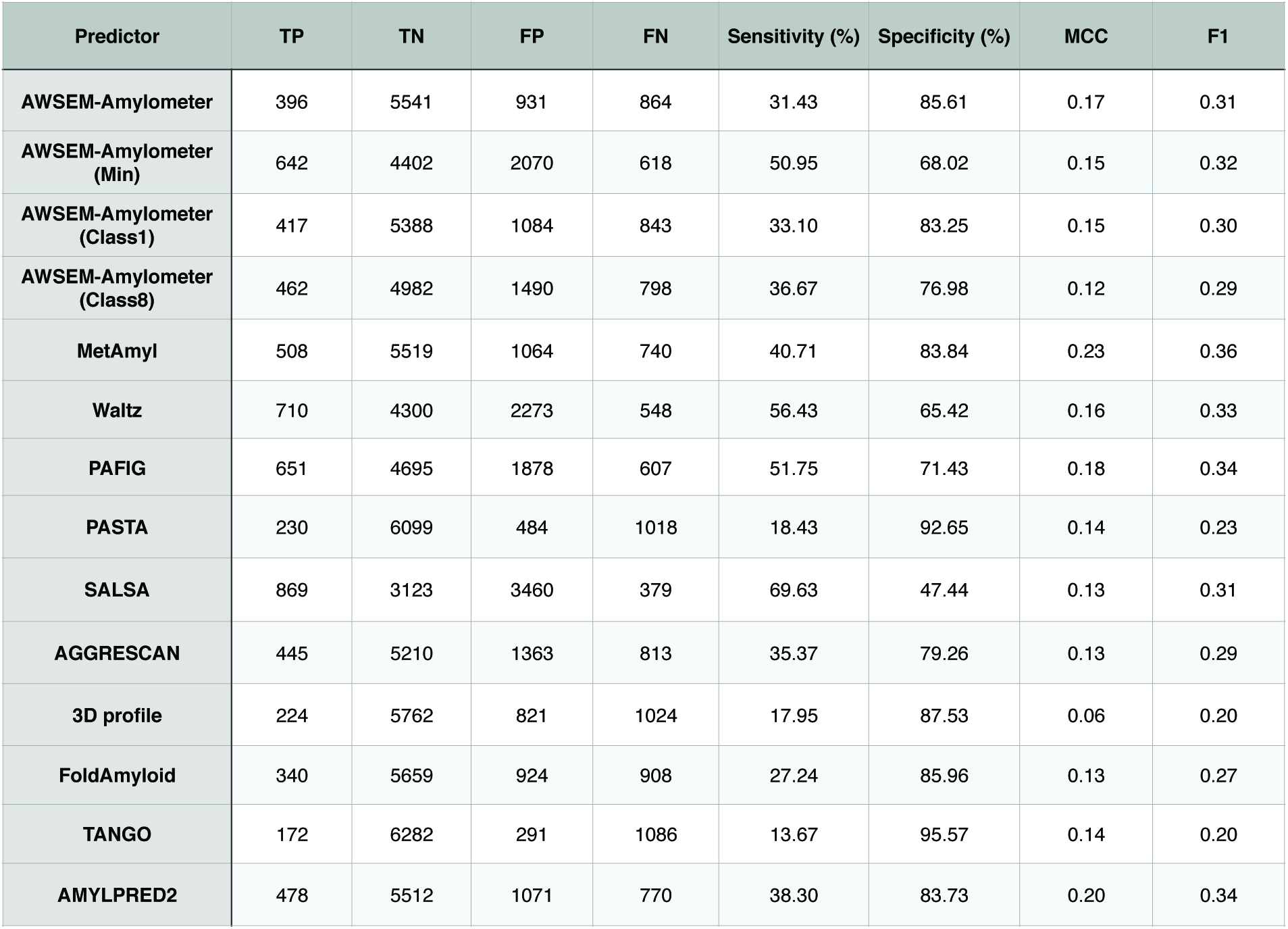
Comparison of prediction performance of the AWSEM-Amylometer on 33 proteins from the amylome with 10 other predictors.

The AWSEM-Amylometer has a lower sensitivity (31.43%) for finding amyloidogenic segments in the amylome dataset of full length proteins compared to its performance for the Waltz dataset of short peptides (62.14%), but it displays a comparable specificity. The Waltz and TANGO algorithms show similar trends. Most of the 33 proteins experimentally studied do not have structures determined for the amyloid fiber to confirm the exact amyloidogenic regions, but the predictions are reasonably accurate for those that do possess well-defined structural information (A*β*_42_ protein and *α*-synuclein, details shown later). Obtaining more accurate predictions of cross-*β* fibril formation propensity based only on the local information contained in hexapeptide sequences may be difficult because the sequence context is not considered in locally informed algorithms. There is a clear need for models that are capable of taking the sequence context of amyloidogenic segments into account.

### 3: Favored sequence features for different amyloid topologies

The group of peptides and proteins that form amyloid fibrils is very diverse but not um­ versal.^9^ The propensity of a given short polypeptide to form amyloid fibrils under some thermodynamic condition depends both on amino acid sequence composition and the order of the amino acids. In this section we investigate which amino acid types are favored in which positions for the seven topological classes of cross-,8 amyloid fibers and for the cross-α fiber by generating random sequences and then examining the sequence preferences for the most stable sequences in each topology.

To find out which amino acids are favored at each position for each of the seven topologies, we generated 5000 random hexapeptide sequences and computed their energies in the seven template structures. For the cross-*α* topology, we generated a different set of 5000 22-residue peptides and computed their energies in the cross-*α* template. Figure 2 shows the energy histograms (left panel) and the sequence logo of the lowest energy peptides (right panel) for each of the eight topologies. For all seven cross-*β* topologies, there are some patterns that are predicted to be most amyloid-prone. The parallel topologies have broader energy distributions. Most amyloid-forming sequence patterns are enriched in Leucines, Isoleucines and Valines, though there are significant differences between the sequence patterns across topologies. In the class 1 topology, the frequency of Glycine and Serine is higher; class 2 and class 8 are dominated by Leucines in the middle of the hexapeptide; class 6 has more charged residues; anti-parallel topology classes 5, 6, and 7 are enriched in valines. Glutamine residues only appear in class 6, meaning that polyglutamine repeats should adopt an anti-parallel topology, a result consistent with published *in vitro* and *in silico* studies. ^27, 32^ Serrano and coworkers have also identified common patterns among amyloidogenic peptides by using mutation scanning experiments.^33^ The *X*_1_*X*_2_*V*_3_*I*_4_*I*_5_*X*_6_ pattern found in their experiments corresponds well with our computed sequence features of the class 7 topology.

**Figure 2:**
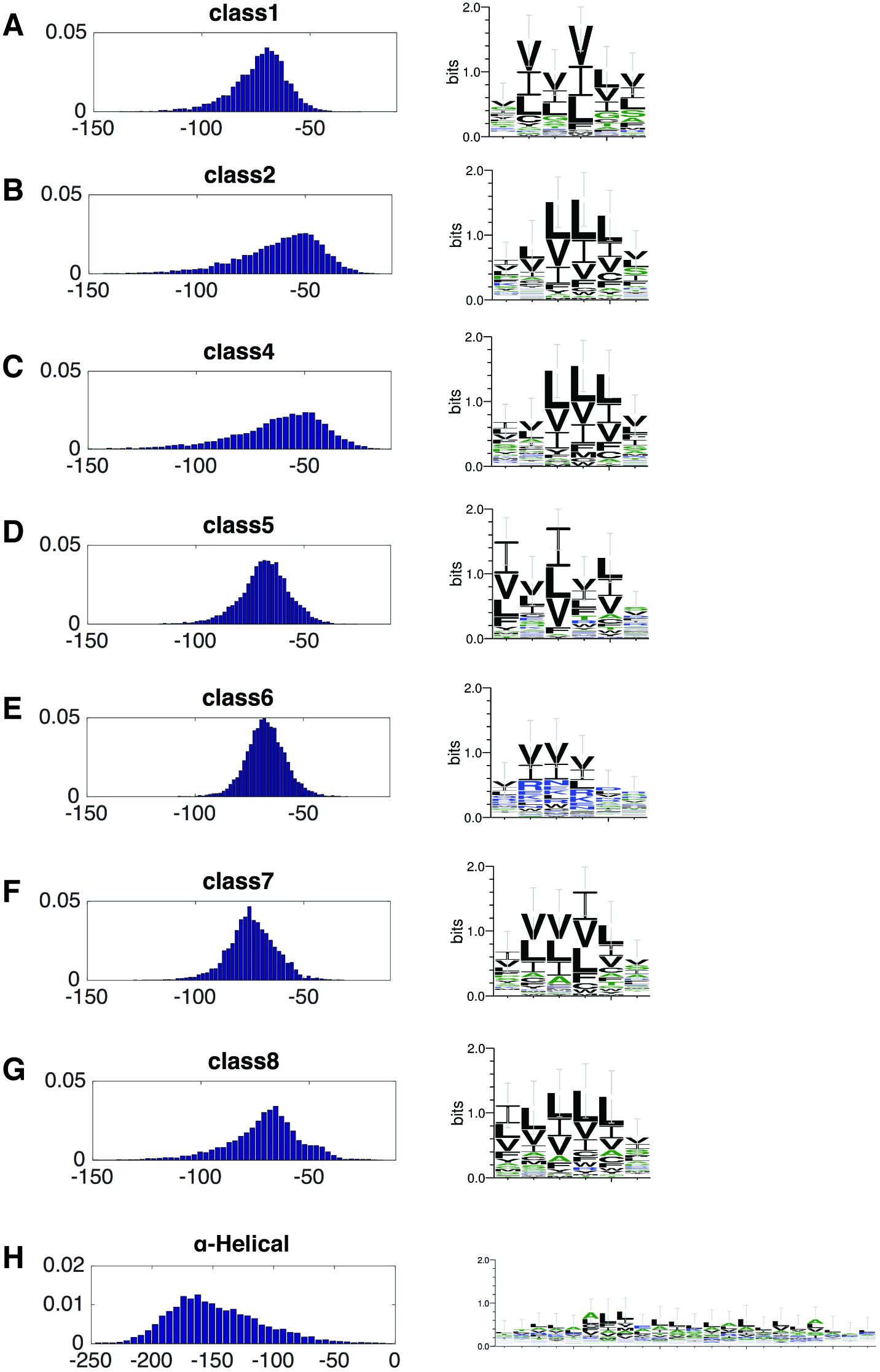
Statistics of the eight topologies on a set of 5000 random sequences. (A) Class 1. (B) Class 2. (C) Class 4. (D) Class 5. (E) Class 6. (F) Class 7. (G) Class 8. (H) cross-α. The left panel is the histogram of the energies for 5000 random sequences and the right panel is the sequence logo for the 50 lowest-energy sequences for each topology.

### 4: The AWSEM-Amylometer is able to predict the topology class of cross-*β* amyloids fibrils

The AWSEM-Amylometer, by evaluating the energy of a hexapeptide sequence in structures with different fibril topologies, not only predicts whether an amyloid should form but also is able to predict the topology of peptide fibrils from sequence data. To check the performance of the AWSEM-Amylometer in predicting fibril topology, we used a dataset of 18 hexapeptides for which well-defined crystal structures have been determined. Among the 18 hexapeptides, the AWSEM-Amylometer successfully predicts the precise topological class of 11 of the peptides, corresponding to an accuracy of 61% (cf. the expected accuracy at random of 1/7 ≈ 14.3%). If we wish only to predict the relative orientation of *β*-strands within a *β*-sheet, we can compare the lowest parallel score (from classes 1, 2 and 4) to the lowest anti-parallel score (from classes 5, 6, 7 and 8). By doing this, the AWSEM-Amylometer predicts correctly the parallel/anti-parallel orientation of 15 peptides (87%, cf. the expected accuracy at random of 1/2 = 50%) (Table 3). For the peptides with apparently incorrect predictions of topology, fibril polymorphism could be contributing to the false negatives. For example, *NNQQNY* is a hexapeptide from the sup35 prion protein of Saccharomyces cerevisiae. An experimentally determined structure shows that the *NNQQNY* peptide has a class 1 cross-*β* structure. In the AWSEM-Amylometer predictions, although *NNQQNY* is predicted to adopt the class 6 anti-parallel topology (with a score of −76.6), its score in the class 1 topology (−67.7) is comparable in value. In keeping with this ambivalence, fibril polymorphism of *NNQQNY* has been found in both *in vitro* and *in silico* studies. ^34, 35^ *AIIGLM* is the A*β*_30–35_ segment from A*β* protein, which forms a parallel *β*-sheet in the crystal structure with PDB ID 2Y3J. The AWSEM-Amylometer predicts that this peptide will adopt an anti-parallel orientation based on a score of −115.36, but a parallel orientation for the segment is predicted to have a nearly equal score of −112.73. The polymorphic tendencies of this peptide have been confirmed by also finding the anti-parallel pattern in the crystal structure of the full length A*β*_40_ segment (PDB ID: 2LNQ). ^36^ While the fragment *KLVFFA*, corresponding to the A*β*_16–21_ segment, adopts an anti-parallel topology in all available hexapeptide crystal structures (PDB ID: 3OW9, 2Y2A and 2Y29), the AWSEM-Amylometer predicts that this hexapeptide is very amyloidogenic (parallel score: −133.72; antiparallel score: −112.53), and the determined structures of *KLVFFA* within the full length A*β*_40_ indeed show both parallel (PDB ID: 2LMQ, 2LMP, 2LMN, 2M4J, 2BEG and 2MXU) and anti-parallel topologies (PDB ID: 2LNQ).

**Table 3:** Topology prediction performance of AWSEM-Amylometer on 18 hexapeptide fiber structures.

| PDB ID | Sequence | Topology Class | Energy Class1 | Energy Class2 | Energy Class4 | Energy Class5 | Energy Class6 | Energy Class7 | Energy Class8 | Predicted Class | Predicted Parallism |
| --- | --- | --- | --- | --- | --- | --- | --- | --- | --- | --- | --- |
| 1YJO | NNQQNY | Class 1 | -67.7 | -52.6 | -49.3 | -72.2 | **-76.6** | -75.6 | -67.4 | Class 6 | **AP** |
| 3FVA | SSTNVG | Class 1 | **-74.8** | -56.6 | -53.9 | -70.4 | -71.2 | -71.4 | -63.4 | Class 1 | **P** |
| 4R0P | IFQINS | Class 1 | **-85.3** | -73.5 | -67.0 | -77.6 | -77.0 | -81.0 | -73.8 | Class 1 | **P** |
| 3NVF | IIHFGS | Class 1 | **-96.90** | -90.44 | -86.29 | -80.52 | -78.15 | -91.17 | -94.54 | Class 1 | **P** |
| 3NVG | MIHFGN | Class 1 | **-89.83** | -76.12 | -79.27 | -73.89 | -71.68 | -87.37 | -87.07 | Class 1 | **P** |
| 3PPD | GGVLVN | Class 1 | **-96.85** | -95.38 | **-100.07** | -79.43 | -73.80 | -89.87 | -95.28 | Class 1 | **P** |
| 2Y3J | AIIGLM | Class 2 | -103.49 | **-108.82** | **-112.73** | -92.28 | -80.48 | -99.66 | **-115.36** | Class 8 | **AP** |
| 5E5X | ANFLVH | Class 2 | -91.37 | -93.98 | **-103.33** | -79.20 | -66.04 | -78.93 | -100.20 | Class 4 | **P** |
| 2ONV | GGVVIA | Class 4 | -105.17 | -108.44 | **-115.29** | -80.82 | -78.24 | -93.13 | -100.73 | Class 4 | **P** |
| 3LOZ | LSFSKD | Class 5 | -67.24 | -48.69 | -41.62 | **-74.82** | -68.02 | -65.74 | -60.65 | Class 5 | **AP** |
| 3NVE | MMHFGN | Class 6 | -72.00 | -61.37 | -73.58 | -62.57 | -55.32 | -80.94 | **-84.16** | Class 8 | **AP** |
| 3OW9 | KLVFFA | Class 7 | -105.19 | **-122.02** | **-133.72** | -76.37 | -71.49 | -97.91 | **-112.53** | Class 4 | **P** |
| 2OMP | LYQLEN | Class 7 | -66.99 | -53.51 | -52.53 | -55.40 | -53.56 | **-69.27** | -69.12 | Class 7 | **AP** |
| 2OMQ | VEALYL | Class 7 | -85.84 | -90.27 | -98.01 | -69.87 | -62.37 | -86.43 | **-100.50** | Class 8 | **AP** |
| 3FR1 | NFLVHS | Class 7 | -90.85 | -94.05 | -94.52 | -81.83 | -76.30 | -92.53 | **-100.06** | Class 8 | **AP** |
| 3NHC | GYMLGS | Class 8 | -80.79 | -80.19 | -86.52 | -67.47 | -64.46 | -77.19 | **-92.87** | Class 8 | **AP** |
| 3NHD | GYVLGS | Class 8 | -88.40 | -86.07 | -89.27 | -75.98 | -73.35 | -89.80 | **-97.69** | Class 8 | **AP** |
| 2ONA | MVGGVV | Class 8 | -97.09 | -86.01 | -91.68 | -79.07 | -74.12 | -93.00 | **-99.09** | Class 8 | **AP** |

We also tested the ability of the AWSEM-Amylometer to predict the relative orientation of *β*-strands within a *β*-sheet on another set of 11 longer peptides where only the parallel/anti-parallel information was available from experiments. As shown in Table 4, the AWSEM-Amylometer successfully predicts the orientation even when experimental evidence suggests ambiguity in the preferred orientation (e.g., the peptide *YTIAALLSPYS* has both parallel and antiparallel topologies in crystal structures and the AWSEM-Amylometer scores both of these configurations as being amyloid prone). In addition to the above peptides exhibiting polymorphism, we also analyzed the conformational preferences of several poly-amino acid peptides and compared the results of the AWSEM-Amylometer to the published information that was available. Polyalanine (*A*_6_) is a common motif in silk fiber, which self-assembles to form antiparallel *β*-sheets. ^37, 38^ The AWSEM-Amylometer predicts that this peptide should adopt an anti-parallel conformation. Polyglutamine repeats are involved in the onset of at least nine neurodegenerative diseases. Fiber structures of polyglutamine repeats show that they prefer an anti-parallel orientation. ^27, 32^ Polyasparagine (*N*_6_) is present in multiple prion-like proteins (e.g., sup35). Simulations carried out by Lindquist and coworkers suggest that this peptide assumes an antiparallel conformation. ^39^ Polyglutamic acid (*E*_6_) forms anti-parallel *β*-sheet according to FTIR experiments. ^39^ All of these results are in keeping with the AWSEM-Amylometer predictions.

**Table 4:** Parallelism prediction performance of AWSEM-Amylometer on 11 short peptide sequences.

| Sequence ID | Sequence | Experimental Topology | Lowest Parallel | Lowest Antiparallel | Predicted Parallelism | Data Source |
| --- | --- | --- | --- | --- | --- | --- |
| **2M5K** | YTIAALLSPYS | AP | -108.7 | -110.9 | **AP/P** | PDB |
| **2M5M** | YTIAALLSPYS | AP | -108.7 | -110.9 | **AP/P** | PDB |
| **2M5N** | YTIAALLSPYS | P | -108.7 | -110.9 | **AP/P** | PDB |
| **3ZPK** | YTIAALLSPYS | AP | -108.7 | -110.9 | **AP/P** | PDB |
| **2NIE** | VKVKVKVKVPPTK<br>VKVKVKVX | AP | -85.77 | -93.11 | **AP** | PDB |
| **2Y3K** | MVGGVVIA | P | -112.11 | -100.35 | **P** | PDB |
| **2Y3L** | MVGGVVIA | P | -112.11 | -100.35 | **P** | PDB |
| **polyA** | AAAAAA | AP | -88.68 | -90.67 | **AP** | Keten et al |
| **polyQ** | QQQQQQ | AP | -64.06 | -81.34 | **AP** | Buchanan et al |
| **polyN** | NNNNNN | AP | -85.49 | -106.88 | **AP** | Halfmann et al |
| **polyE** | EEEEEE | AP | -54.94 | -73.54 | **AP** | Hernik et al |

The AWSEM-Amylometer is based on the AWSEM force field, which was optimized using principles from energy landscape theory. ^40^ While computational power has been increasing exponentially over the past decades, the complete folding from scratch of even a moderate size protein remains challenging using atomistic force fields. The coarse-grained AWSEM force field has been used to predict the structures of protein monomers and dimers. ^20–23^ We have also recently used the AWSEM force field to simulate and characterize the aggregation of a glutamine-rich mechanical prion CPEB, ^26^ I27, ^25^ A*β*_40_ protein, ^28^ polyglutamine repeats, ^27^ and Huntingtin-Exon-1 encoded protein fragments. ^29^ These studies show that simulations with the AWSEM force field can not only be used to characterize structural changes during protein aggregation efficiently, which are otherwise very difficult to characterize in detail using biophysical techniques, but also to construct aggregation free energy landscapes that are useful for understanding aggregation experiments.

To further demonstrate the capability of the AWSEM-Amylometer in predicting fiber topology, as well as the power of the AWSEM force field to characterize further the aggregation process efficiently, we used molecular dynamics simulations with AWSEM to construct the aggregation free energy landscapes of three different hexapeptides (*GGVVIA*, *GYMLGS* and *Q*_6_). The number of parallel hydrogen bonds and antiparallel hydrogen bonds were used to evaluate the topology of the simulated fiber structures. Figure 3 shows free energy landscapes of the three hexapeptides (*GGVVIA*, *GYMLGS* and *Q*_6_) computed with AWSEM. The preference for these hexapeptides to adopt parallel versus anti-parallel topologies is reflected in the free energy minima on the computed free energy landscapes. These minima correspond to the experimentally determined preferences for all three of the peptides.

**Figure 3:**
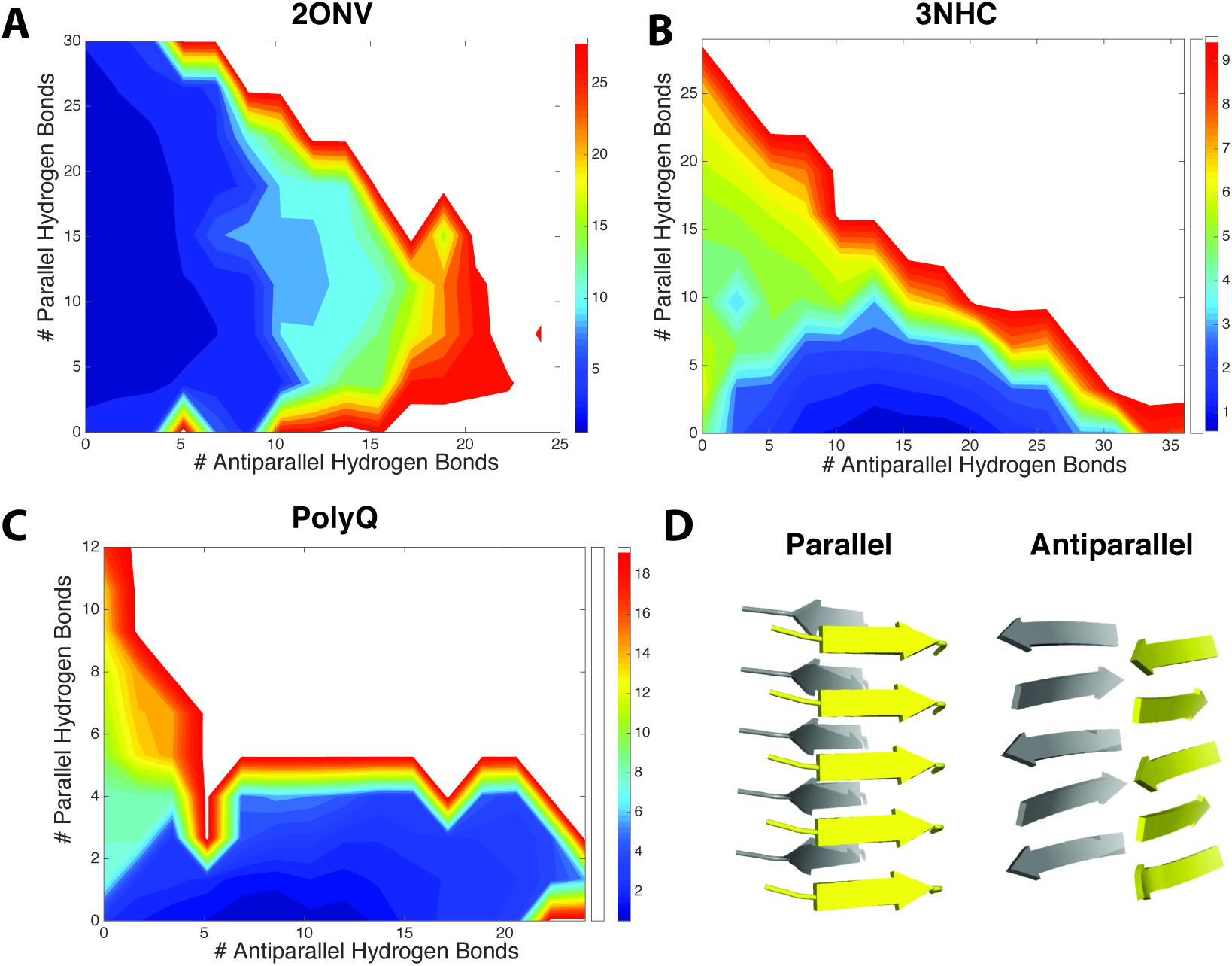
Free energy landscapes of hexapeptides. The number of anti-parallel hydrogen bonds and the number of parallel hydrogen bonds are used as order parameters to show if the peptides prefer to adopt a parallel or anti-parallel topology. The free energy surfaces of (A) *GGVVIA*, which favors a parallel topology, (B) *GYMLGS*, which favors an antiparallel topology, and (C) *Q*_6_, which favors an anti-parallel topology. Examples of parallel and anti-parallel topologies are shown in (D).

### 5: Prediction ambiguity and amyloid polymorphism of *β* steric zippers

Amyloid fibril polymorphism has multiple causes, including the sequence context and the solvent conditions. Fibril topology may not be exclusively determined by the local sequence. One of the most intriguing features of sequence-encoded polymorphism is that the same peptide can adopt distinct chain-folding patterns that give rise to a variety of cross-*β* structures. ^41, 42^ This type of polymorphism can lead to different amyloid strains. There is often a barrier of propagation or transmission between different strains (e.g. in Sup35, A*β*), which makes cross seeding impossible or at least inefficient. Understanding how polymorphism is encoded by protein sequence is key to understanding the species barriers that arise from these molecular-level structural details. In addition to the predicted polymorphisms in the peptides mentioned above, the AWSEM-Amylometer is also able to predict the possibility of polymorphism for longer protein sequences and, therefore, should be useful in predicting species barriers.

Amyloid polymorphism for A*β* has been studied extensively by Eisenberg and coworkers. These studies have not pinpointed why A*β* can assume both parallel and anti-parallel orientations. ^36^ In the case of A*β* fibers, the AWSEM-Amylometer suggests that A*β* can adopt both parallel and anti-parallel conformations (Cyan lines in Figure 4 A, B). As shown in the previous section, A*β*_16–21_ and A*β*_30–35_, the two core-regions for fiber formation as revealed by crystal structures, both demonstrate strong ambiguity in their preferred orientation, thus leading to the polymorphism in full-length fiber structures. Zheng et al. demonstrated that there is a profound change in amyloidogenicity even from point mutations using only the *NNQQNY* topology. ^28^ Our results show that these point mutations can generate similar changes in an antiparallel topology (Figure 4B): increased hydrophobicity at site 22 elevates the amyloidogenicity of the hexapeptides that contain this site, and *E*22*V* is more amyloidogenic compared to *E*22*G* and *E*22*Q* in both parallel topology and antiparallel topology. Similarly, the AWSEM-Amylometer predicts that *α*-synuclein should exhibit both parallel and anti-parallel structures (Figure S1). This result is consistent with the diversity of experimental results that have been reported regarding the relative orientation of *β*-strands within *α*-synuclein fibers. ^43, 44^

**Figure 4:**
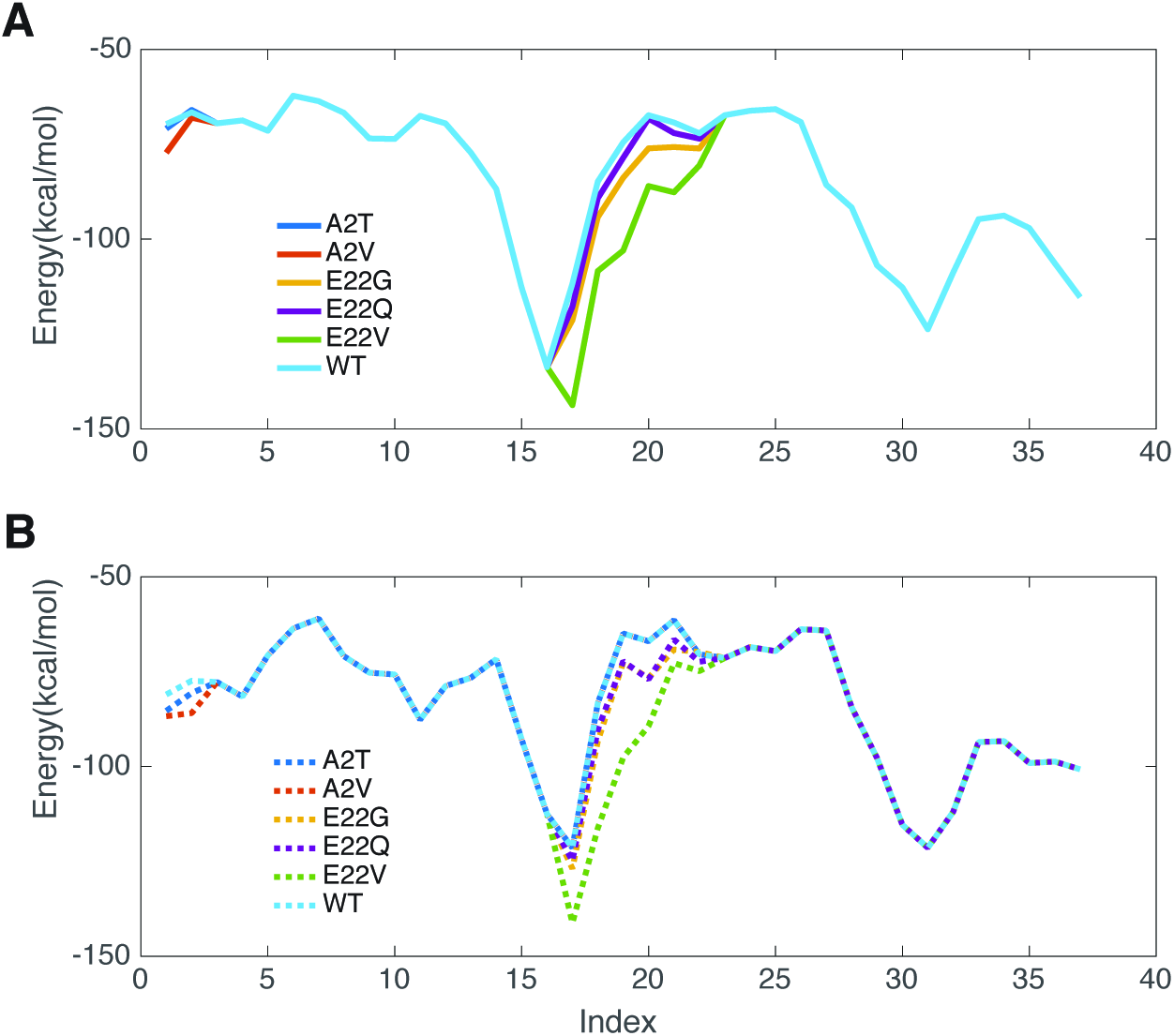
Calculated propensity of A*β*_42_ to form parallel (A) and anti-parallel (B) steric zippers.

One has to admit that a model that is only locally informed and that focuses on hexapeptides by themselves must be limited in its capability to elucidate the topology of amyloids formed by full-length proteins. The problem of locality intrinsic to the AWSEM-Amylometer is shared by other predictors. The connection to the AWSEM force field, which is optimized for globular protein folding and native structure prediction, however, has enabled us to go beyond the purely local characterization to conduct direct protein aggregation studies *in silico*, such as those we have carried out for the aggregation of I27, ^10^ A*β*_40_, ^28^ polyglutamine repeats ^27^ and HTT-exon1 encoded protein fragments. ^29^ Simulations using the AWSEM force field not only capture the local signatures seen from the AWSEM-Amylometer calculations (i.e. that A*β*_40_ could have both parallel and antiparallel structures in simulations ^28^), but also allow one to find the most favorable structure that lies in the “amyloid funnel”.^27, 28^

### 6: The AWSEM-Amylometer can be used to predict the propensity for forming *α*-helical amyloid fibers

Previously, the cross-*β* spine, in which stacked *β*-strands run perpendicular to the fibril axis, was believed to be the universal architecture for all amyloid structures that bind thioflavin T.^2^ Tayeb-Fligelman et al. recently showed that phenol-soluble modulin *α*-3 (PSM*α*3) whose aggregates pass the standard laboratory amyloid criteria (e.g. binding thioflavin T) actually forms amphipathic *α*-helices that pack together to form an unusual cross-*α* fiber topology. ^45^ In order to understand the propensity of this 22-residue peptide to form a cross-*α* fiber and compare it to other sequences without experimentally solved fiber structures, we used the AWSEM-Amylometer to compute the energy of the PSM*α*3 sequence and related sequences when taking on this structure. The PSM*α*3 sequence is highly favored in the cross-*α* topology (score: −214.49) compared to the distribution of energies for 5000 random sequences threaded on this topology (Figure 2H). According to the cross-*β* AWSEM-Amylometer, PSM*α*3 is unlikely to form cross-*β* fibers (Figure 5A). PSM*α*3 mutants *K* 9*P /F* 11*P* and *F* 3*A* significantly reduce fiber formation and its related toxicity according to experiments, while the *G*16*A* mutation was found to enhance toxicity. ^45^ The AWSEM-Amylometer calculations for these mutants correspond well with these experiments in that they show the *K* 9*P /F* 11*P* and *F* 3*A* variants have higher energies than the wild type (less favorable in the cross-*α* template), while the *G*16*A* mutation significantly lowers the energy. The AWSEM-Amylometer suggests that other PSMs like PSM*α*1, PSM*α*2 and PSM*α*4, are also very likely to form cross-*α* amyloids (Table 5). The PSM*β*1-2 peptides show relatively weak signals, indicating that, if the PSM*β*1-2 peptides form cross-*α* fibers, the core structure may be somewhat different from that of PSM*α*3.

**Figure 5:**
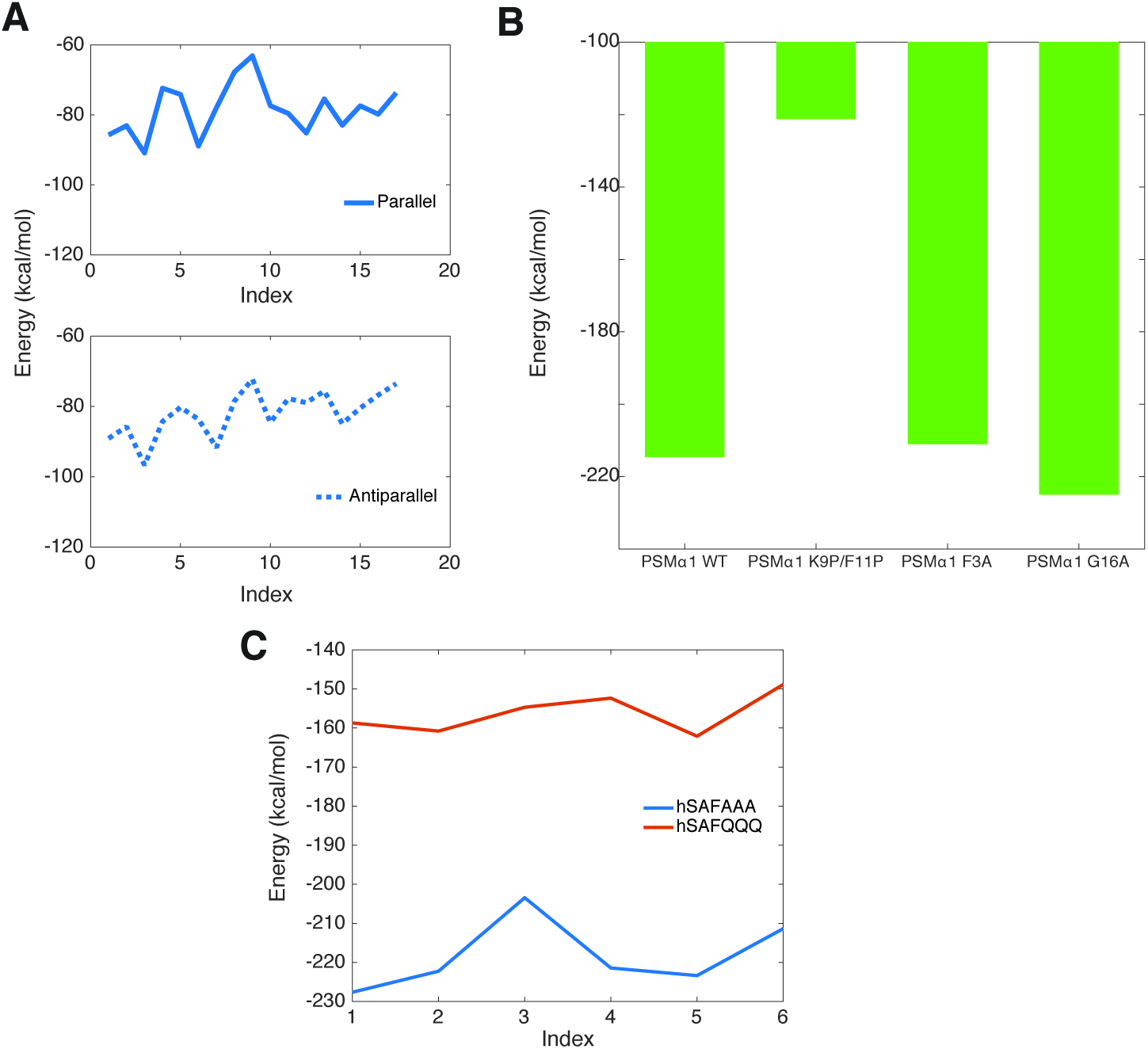
Calculated propensity to form cross-*β* and cross-*α* fibrils for PSMs and synthetic peptides (hSAFAAA and hSAFQQQ). A: Calculated propensity of PSM*α*-3 to form parallel (upper, solid line) and anti-parallel (lower, dashed line) *β*-sheet structures. B: Calculated propensity of PSM*α*-3 and its mutants to form *α*-helical amyloid. The F3A mutant and the K9P/F11P double mutant, which do not form fibrils, exhibited lower propensity to form cross-*α* fibers compared to wildtype PSM*α*-3, while G16A is predicted to have a higher propensity to form cross-*α* fibers. C: Calculated propensity of the synthetic peptides *hSAFAAA* (red line) and *hSAFQQQ* (blue line) to form cross-*α* fibers.

**Table 5:**
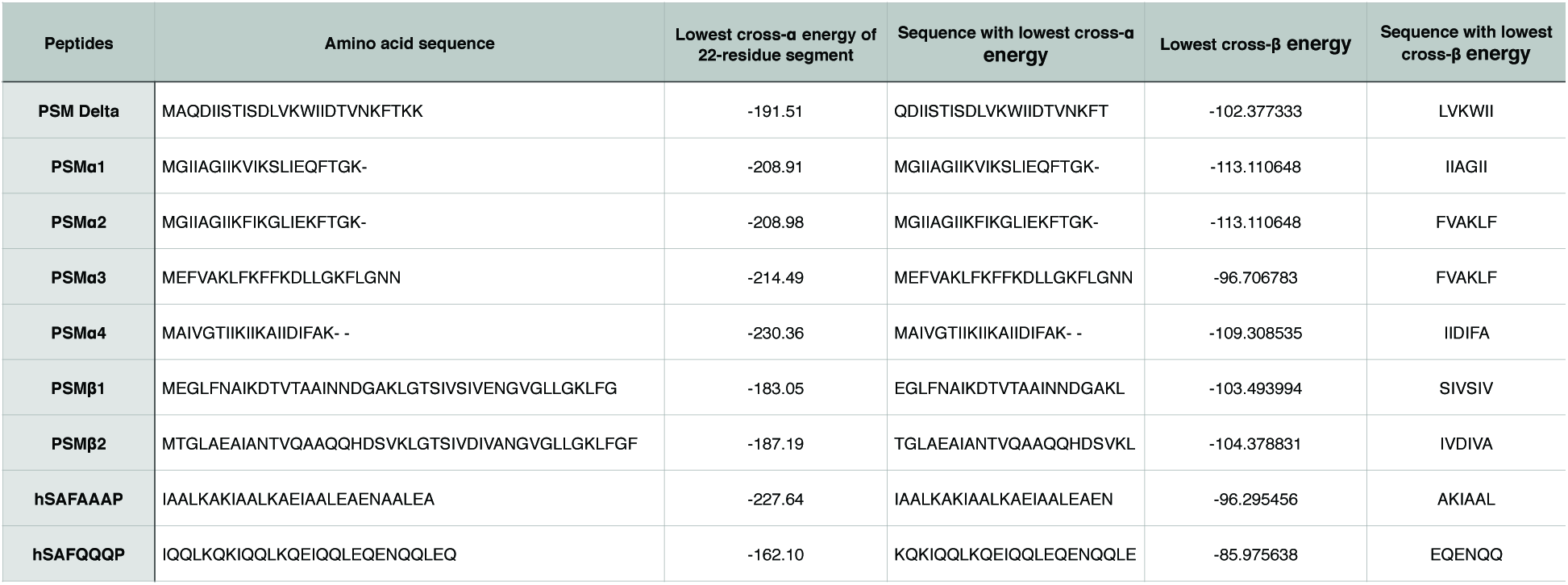
List of sequences and AWSEM-Amylometer scores for 9 proteins in the cross-*α* topology.

In addition to the naturally occurring PSMs, synthetic systems have been found that form *α*-helical fibrils. Banwell et al. have used the synthetic peptide *hSAF AAA* to form hydrogels that contain *α*-helical fibrils. ^46^ *hSAF AAA* turns out to be very amyloid-prone in the cross-*α* topology according to the AWSEM-Amylometer, while another peptide, *hSAF QQQ*, does not favor this topology (Figure 5C). This result agrees well with the experiments showing that *hSAF QQQ* formed *β*-sheet containing structures. ^46^

## Conclusion

In conclusion, we have explored the power of the AWSEM-Amylometer to scan for the amyloidogenic segments and assign their topologies in the fibers that form. The present study on the Waltz dataset of peptides documents the prediction capabilities of the AWSEM-Amylometer for peptides. In contrast to other predictors, the AWSEM-Amylometer also provides accurate predictions of the topologies of amyloids. Simulations and structure predictions using the AWSEM force field can be used to further characterize the topological preferences efficiently for multiple hexapeptides. As we have evidenced in previous work on I27, A*β*, polyglutamine repeats and HTT-exon1 encoded protein fragments, the AWSEM force field can also capture nonlocal effects that go beyond the reach of other locally informed prediction approaches.

## Acknowledgement

This work was supported by Grant R01 GM44557 from the National Institute of General Medical Sciences. Additional support was also provided by the D.R. Bullard-Welch Chair at Rice University, Grant C-0016. We thank the Data Analysis and Visualization Cyberinfrastructure funded by National Science Foundation Grant OCI-0959097.

